# Beyond antigen delivery: tailoring the Coated Bacterial Vaccine platform with a *Lentilactobacillus kefiri* S-layer protein to modulate vaccine-induced immunity

**DOI:** 10.64898/2026.11.24.768567

**Authors:** Inés Harguindeguy, Matías H. Assandri, Melisa S. Homse, María A. Daza Millone, María de los A. Serradell, Gastón E. Ortiz

## Abstract

Coated Bacterial Vaccines (CBVs) are a versatile antigen display platform based on chemically inactivated *Bacillus subtilis*. We previously showed that CBVs induce protective immune responses with a predominant Th2 profile and are effective with both strongly and weakly immunogenic antigens. Here, we explored whether the S-layer protein SLP 83111 from *Lentilactobacillus kefiri* can be co-immobilized on CBVs to modulate this response. Native SLP 83111 was co-immobilized with the C-terminal fragment of tetanus toxin (TTFC-dSLPA) on inactivated *B. subtilis*, and binding was characterized by MP-SPR and adsorption isotherms. BALB/c mice were immunized with CBVs-TTFC, CBVs-TTFC-SLP, or soluble TTFC, and anti-TTFC IgG, IgG1 and IgG2a levels, protection against lethal tetanus toxin challenge and cytokine expression in RAW 264.7 macrophages were evaluated. SLP 83111 bound to the carrier with high affinity (K_D_∼10^-8^ M; B_max_ 25 µg/OD), and its co-incorporation with TTFC was confirmed by SDS-PAGE and Western blot. CBVs-TTFC induced a strongly Th2-biased response (IgG1/IgG2a ratio 30:1), whereas CBVs-TTFC-SLP displayed a more balanced ratio (5:1), mainly due to lower IgG1 titers. CBVs-TTFC-SLP showed a trend toward improved survival compared with CBVs-TTFC and significantly outperformed soluble TTFC. In macrophages, SLP 83111 reduced IL-1β expression without affecting IL-6 or IL-10. These results demonstrate that the CBVs platform can be rationally modulated, extending its scope beyond antigen delivery and supporting the design of tailored bacterial vaccines.

## 1. Introduction

Vaccines have been one of the most effective interventions in the history of medicine, drastically reducing morbidity and mortality from infectious diseases worldwide [1–3]. Their evolution, from live-attenuated microorganisms to mRNA-based and vectored vaccines, reflects a continuous search for safer, more versatile and more efficacious immunization strategies [4, 5]. Beyond the selection of protective antigens, the rational design of vaccine formulations requires delivery systems capable of promoting immune responses suited to the target pathogen or disease. In this context, particulate antigen delivery platforms have attracted considerable attention because they enhance antigen presentation, facilitate interactions with antigen-presenting cells (APCs), and provide immunostimulatory signals that contribute to the activation of innate and adaptive immunity. Nevertheless, modulating the qualitative characteristics of the immune response remains a major challenge in vaccine development.

Bacterial-based antigen delivery systems are particularly attractive because their cell envelopes contain pathogen-associated molecular patterns (PAMPs) that are recognized by pattern recognition receptors (PRRs) on innate immune cells, contributing to APC activation and shaping the subsequent adaptive response [6, 7]. However, the immunological profile elicited by a given bacterial carrier may not be optimal for every antigen or vaccination objective. Live bacterial vaccines (LBVs) and bacterium-like particles (BLPs) have been extensively explored, but each presents limitations: LBVs rely on genetically modified organisms, which poses regulatory obstacles [8, 9], whereas BLPs require acid and thermal treatments that degrade immunostimulatory components such as bacterial DNA and surface proteins [10, 11]. Decorated live bacteria have also been proposed, but the recombinant antigen tends to dilute upon cell division [12]. These constraints highlight the need for antigen display systems that preserve the structural and immunological integrity of whole bacteria while avoiding genetic modification and antigen dilution.

Recently, we developed the Coated Bacterial Vaccines (CBVs) platform, a non-genetically modified delivery system based on chemically inactivated *Bacillus subtilis* cells coated with recombinant antigens through non-covalent interactions [13]. Antigen immobilization is mediated by a protein anchoring domain derived from the S-layer protein SlpA of *Lactobacillus helveticus*, allowing recombinant proteins to associate with the bacterial surface without requiring genetic modification of the carrier. Using the C-terminal fragment of tetanus toxin (TTFC) as a model antigen, CBVs induced antigen-specific antibody responses and conferred full protection against lethal tetanus toxin challenge in mice [13].

The versatility of the platform was subsequently explored with a recombinant antigen containing repeated gonadotropin-releasing hormone (GnRH) sequences, a weakly immunogenic target relevant to immunocastration strategies [14]. Although CBVs elicited functional antigen-specific responses in murine and porcine models, the immune responses were characterized by a predominance of IgG1 over IgG2a, consistent with a Th2-associated humoral profile [13, 14]. This profile was effective in the models evaluated, but the ability to modify the immunological properties of the platform could broaden its applicability to antigens requiring different response profiles. Notably, the intrinsic PAMPs displayed on the *B. subtilis* surface appear to be the main drivers of this Th2 bias, suggesting that the incorporation of additional immunomodulatory components could redirect or balance the response.

Bacterial S-layer proteins (SLPs) are particularly interesting in this regard. These proteins form highly organized surface layers in numerous bacterial species and display structural and biochemical properties that influence their interactions with host cells. Several SLPs from lactic acid bacteria have been shown to interact with immune cells and modulate inflammatory and immunoregulatory signaling pathways, with activities that vary substantially among proteins from different species and strains. In particular, SLPs from *Lentilactobacillus kefiri* have been investigated for their ability to interact with APCs. SLP 8348 is internalized by murine RAW 264.7 macrophages through carbohydrate-dependent interactions and enhances LPS-induced activation, including the expression of antigen-presentation and costimulatory molecules and the production of IL-6 and IL-10 [15]. SLP 8321 and SLP 5818 activate dendritic cells through C-type lectin receptors such as Mincle and SignR3, respectively, promoting cytokine production and costimulatory molecule expression [16, 17].

Among these proteins, the native SLP from *L. kefiri* CIDCA 83111 (SLP 83111) is particularly attractive as a potential immunomodulatory component. SLP 83111 has been reported to enhance APC activation on its own and to exert adjuvant-like effects on antigen-specific antibody responses when co-administered with S-layer proteins from *Clostridioides difficile* [18]. These findings suggest that SLP 83111 could provide additional immunological signals when incorporated into a particulate antigen delivery system.

Based on these observations, we hypothesized that native SLP 83111 could be co-immobilized on CBVs and shift the anti-TTFC response toward a more balanced IgG1/IgG2a ratio without modifying the antigen or the carrier. We characterized SLP 83111 binding to inactivated *B. subtilis*, established sequential co-immobilization with TTFC-dSLPA, and evaluated IgG and IgG subclass responses, protection against lethal tetanus toxin challenge, and cytokine expression in murine macrophages. The central aim of this work was to demonstrate that the CBVs platform itself can be rationally modulated, extending its scope beyond antigen delivery and supporting the design of tailored bacterial vaccines.

## 2. Materials and methods

### 2.1 Bacterial strains and culture conditions

*Escherichia coli* BL21-CodonPlus (DE3)-RIPL was used for heterologous protein expression and was grown in Luria-Bertani (LB) broth or LB-agar plates at 37 °C with shaking at 200 rpm, as previously described [13]. *Bacillus subtilis* var. natto (ATCC 15245) was used as the antigen-presenting carrier and was cultured under the same conditions [13]. *Lentilactobacillus kefiri* CIDCA 83111 was grown in de Man, Rogosa and Sharpe (MRS) broth (Difco) under aerobic conditions at 37 °C for 48 h, without agitation. Bacterial growth was monitored by measuring the optical density at 600 nm (OD_600_).

### 2.2 Production of the recombinant antigen TTFC-dSLPA

The recombinant antigen TTFC-dSLPA, consisting of the C-terminal fragment of tetanus toxin (TTFC) fused to the carboxy-terminal cell-wall-binding domain of the SlpA protein from *Lactobacillus helveticu*s (dSLPA), was produced in *E. coli* BL21 transformed with the pET-TTFC-dSLPA plasmid, as described in detail previously [13]. Briefly, cultures were induced with 1 mM IPTG at 20 °C for 8 h, cells were lysed by sonication, and the soluble fraction was recovered by centrifugation. The antigen was purified by immobilization on inactivated *B. subtilis* followed by elution with 100 mM carbonate-bicarbonate buffer pH 10.2, as reported [13, 22]. Purified TTFC-dSLPA was dialyzed against PBS, quantified by BCA assay (Pierce, Thermo Fisher), and its integrity was verified by SDS-PAGE.

### 2.3 Preparation of the bacterial carrier

The bacterial carrier was prepared from *B. subtilis* var. natto (ATCC 15245) as previously described [13]. Briefly, an overnight culture was harvested by centrifugation at 5,000 x g for 5 min at 4 °C, washed three times with PBS, and inactivated by incubation with 2% (v/v) glutaraldehyde for 16 h at room temperature in a rotator. Inactivated cells were washed three times with PBS, counted in a Neubauer chamber, and adjusted to 2×10^9^ cells/mL.

### 2.4 Extraction of native SLP 83111

Native SLP 83111 was extracted from the surface of *L. kefiri* CIDCA 83111 following the protocol of Malamud and co-workers [15]. Briefly, stationary-phase cells were harvested by centrifugation at 10,000 x g for 5 min at 10 °C, washed twice with sterile PBS, and resuspended in 5 M LiCl. The suspension was vortexed for 10 min and centrifuged at 12,000 x g for 15 min at 10 °C. The supernatant was filtered through a 0.45 µm membrane and dialyzed against PBS for 36 h at room temperature with two buffer changes, using cellulose tubing (12.4 kDa cutoff, Sigma-Aldrich). The dialysate was centrifuged at 12,000 x g for 15 min at 4 °C and the protein concentration was determined using the Pierce BCA Protein Assay Kit (Thermo Fisher). The integrity and purity of SLP 83111 were verified by 12% SDS-PAGE.

### 2.5 Formulation of tailored CBVs with TTFC and SLP

Tailored CBVs formulations were prepared by binding the proteins of interest to the surface of the bacterial carrier, following the general procedure described previously [13]. For formulations incorporating SLP 83111, a sequential coating strategy was used: the carrier was first incubated with TTFC-dSLPA and then with SLP 83111. Briefly, 60 OD_600_ units of the bacterial carrier were incubated with 1,500 µL of TTFC-dSLPA (1.8 µM) at 80 rpm for 30 min at room temperature. The pellet was washed three times with PBS and divided into two fractions. One fraction was kept as CBVs-TTFC, and the other was incubated with 300 µL of SLP 83111 (1.4 µM) to produce CBVs-TTFC-SLP. CBVs were centrifuged at 5,000 x g for 10 min at 4 °C, washed three times with PBS, and finally resuspended in PBS. Protein concentration was determined by SDS-PAGE followed by densitometric analysis, and the co-incorporation of both proteins on the carrier was verified by SDS-PAGE and Western blot.

### 2.6 SDS-PAGE and Western blot

Samples were resolved by 12% SDS-PAGE and stained with Coomassie brilliant blue, as described [13]. For Western blot, proteins were transferred to a nitrocellulose membrane by electrotransfer (300 mA, 60 min) in Tris-glycine buffer containing 20% (v/v) methanol using a Mini Trans-Blot System (Bio-Rad). Membranes were blocked with 3% (w/v) skim milk in PBS for 1 h at 37 °C and incubated with mouse polyclonal anti-TTFC or anti-SLP 83111 sera for 2 h at 37 °C. After washing, membranes were incubated with horseradish peroxidase-conjugated anti-mouse IgG (1:2,000, Sigma-Aldrich) for 1 h at 37 °C. Detection was performed using 4-chloro-1-naphthol (0.5 mg/mL) in TBS containing 17% (v/v) methanol and 0.15% (v/v) H₂O₂. Purified TTFC-dSLPA and SLP 83111 were included as controls. Band identity was confirmed by comparison with controls and with the expected molecular weights (TTFC-dSLPA: ∼72 kDa; native SLP 83111: ∼71 kDa). Polyclonal anti-TTFC serum was obtained from mice immunized with purified TTFC-dSLPA according to conventional immunization protocols [24]; polyclonal anti-SLP 83111 serum was obtained following the same procedure using purified native SLP 83111 as immunogen.

### 2.7 Multi-parametric surface plasmon resonance (MP-SPR)

The affinity and binding kinetics of TTFC-dSLPA and SLP 83111 to the bacterial carrier were evaluated by MP-SPR, following the protocol described in detail previously [14]. A gold sensor surface treated with 0.01% (w/v) poly-L-lysine was coated with 100 µL of a 2 OD_600_/mL suspension of inactivated *B. subtilis*. The surface was then blocked with 500 µL of BSA 5 mg/mL, washed with Milli-Q water, and dried under a nitrogen stream. All incubations were performed for 1 h at room temperature. Increasing concentrations of purified TTFC-dSLPA (5, 10 and 15 µg/mL) and SLP 83111 (1.3, 2.6 and 4.5 µg/mL) in PBS–Tween 20 0.05% (v/v) were injected at a flow rate of 10 µL/min for 10 min, followed by a 10-min wash. Sensorgrams were analyzed with TraceDrawer v1.5 using a one-to-one interaction model. Association (k□) and dissociation (k_d_) rate constants and the equilibrium dissociation constant (K_D_) were calculated from independent fits of the association and dissociation phases (n = 3 independent experiments, each with four technical replicates).

### 2.8 Adsorption isotherm for SLP 83111

The maximum binding capacity (B_max_) and apparent dissociation constant (K_d_) of SLP 83111 were determined by a saturation assay, as described previously [13]. Briefly, a fixed amount of bacterial carrier (1×10^8^ cells) was incubated with increasing amounts of purified SLP 83111 (0.3-3 µM) for 30 min at 25 °C. After incubation, cells were centrifuged and washed three times with PBS, and the amount of protein bound to the bacterial surface was quantified by SDS-PAGE and densitometric analysis, using purified SLP 83111 as standard. The isotherm was fitted to a sigmoidal model to obtain B_max_ and K_d_ using GraphPad Prism 6.0 (GraphPad Software), as described [13].

### 2.9 Immunization and tetanus toxin challenge in mice

Female BALB/c mice (n = 5 per group), 6–8 weeks old, were obtained from the animal facility of the Faculty of Veterinary Sciences, National University of La Plata. Animals were immunized intraperitoneally (i.p.) on days 0, 14 and 28 with 100 µL/dose of the corresponding formulations (Table 1). The CBVs-TTFC group received 5×10^7^ CBVs containing 2.5 µg/dose of TTFC-dSLPA in PBS. The CBVs-TTFC-SLP group received 5×10^7^ CBVs containing 2.5 µg/dose of TTFC-dSLPA and 1 µg/dose of SLP 83111 in PBS. The antigen control group received 2.5 µg/dose of purified TTFC-dSLPA in PBS. Animals were monitored regularly for general condition and adverse reactions. Blood samples were collected at 14-, 28- and 35-days post-immunization (dpi), and sera were stored at −20 °C until analysis. On day 49 post-immunization, mice were challenged i.p. with 75 x LD_50_ of tetanus toxin (TeTx) and monitored for 10 days to evaluate the onset of symptoms, such as muscle rigidity and spasms, and survival rate. TeTx was generously provided by the “Instituto Biológico Dr. Tomás Perón”. All animal procedures were approved by the Institutional Animal Care and Use Committee of the Faculty of Exact Sciences, National University of La Plata (protocol 010-42-24), and performed in accordance with national and international guidelines.

**Table 1.** Composition of the vaccine formulations used in this study. Groups are named according to their antigen and immunomodulator content: “CBVs-TTFC” (TTFC-dSLPA displayed on the carrier), “CBVs-TTFC-SLP” (TTFC-dSLPA and SLP 83111 co-displayed) and “Soluble TTFC” (antigen-only control).

| Group | Formulation | Composition (per dose) |
|---|---|---|
| **1** | CBVs-TTFC | 5x10<sup>7</sup> CBVs displaying 2.5 µg TTFC-dSLPA in PBS |
| **2** | CBVs-TTFC-SLP | 5x10<sup>7</sup> CBVs displaying 2.5 µg TTFC-dSLPA and 1 µg native SLP 83111 in PBS |
| **3** | TTFC | 2.5 µg purified TTFC-dSLPA in PBS (no carrier) |

### 2.10 Determination of anti-TTFC antibodies by ELISA

Levels of total IgG, IgG1 and IgG2a specific for TTFC were determined by indirect ELISA as described previously [13], with minor modifications. Plates (MaxiSorp, Nunc) were coated with 250 ng/well of purified TTFC-dSLPA in 100 mM carbonate-bicarbonate buffer pH 9.6 for 16 h at 4 °C. After blocking with 3% (w/v) skim milk in PBS for 1 h at 30 °C, plates were incubated with 50 µL/well of serial dilutions of sera from immunized animals in 1% (w/v) skim milk in PBS for 1 h at 30 °C. Plates were then incubated with horseradish peroxidase-conjugated anti-mouse total IgG (Sigma-Aldrich), anti-IgG1 or anti-IgG2a (Abcam) for 1 h at 30 °C. The reaction was developed with 2 mg/mL o-phenylenediamine in 0.05 M citrate buffer pH 5.0 containing 0.03% (v/v) H₂O₂, stopped with 2 N H₂SO₄, and absorbance was measured at 492 nm. For group-level comparisons, sera from each group were pooled (5 µL per animal) and antibody titers were calculated as the mid-log_10_ of the reciprocal serum dilution, using the EC_50_ approach [28]. For individual animals, IgG, IgG1 and IgG2a levels are reported as absorbance values at 492 nm at the corresponding cut-off dilutions (1:1,000 for total IgG; dilutions determined for each subclass).

### 2.11 Macrophage stimulation and cytokine mRNA quantification

Murine RAW 264.7 macrophages were cultured and stimulated as described previously [13]. Briefly, cells (4×10^5^ cells/well) were incubated for 6 h with purified SLP 83111 (1 µg/well), CBVs-TTFC (2×10^7^ cells containing 2 µg of TTFC-dSLPA/well), or CBVs-TTFC-SLP (2 × 10 cells containing 2 µg of TTFC-dSLPA and 1 µg of SLP 83111/well). DMEM and lipopolysaccharide (LPS from *E. coli* O111:B4, 2 µg/mL) were used as negative and positive controls, respectively. To minimize residual LPS effects, polymyxin B (10 µg/mL) was added to all conditions except the positive control [26]. After treatment, cells were centrifuged at 3,000 x g for 10 min and pellets were collected for analysis. Total RNA was extracted and reverse-transcribed as described [13, 27], and the relative mRNA expression of IL-6, IL-1β, IL-10 and IL-12 was determined by RT-qPCR using the 2^-ΔΔCt^ method with β-actin as housekeeping gene [27]. Primer sequences were those reported previously [13].

### 2.12 Statistical analysis

All results are expressed as mean ± standard error of the mean (SEM), unless otherwise stated. Statistical analysis was performed with GraphPad Prism 6.0 (GraphPad Software). Significance between two groups was determined by the Mann-Whitney U test. For multiple comparisons, one-way ANOVA was followed by Dunnett’s or Tukey’s post hoc test, as indicated in each figure legend. Survival curves were compared by the log-rank (Mantel-Cox) test. Differences were considered statistically significant at p < 0.05.

## 3. Results

### 3.1 Production of native SLP 83111 and characterization

Native SLP 83111 was successfully extracted from the surface of *L. kefiri* CIDCA 83111 by 5 M LiCl treatment. SDS-PAGE analysis revealed a major band of approximately 71 kDa in the extract, corresponding to the expected molecular weight of SLP 83111, which was obtained with a high degree of purity (Figure 1A). The absence of this band in the bacterial pellet after extraction confirmed the efficient release of the protein from the lactobacillus surface (Figure 1A).

**Figure 1.**
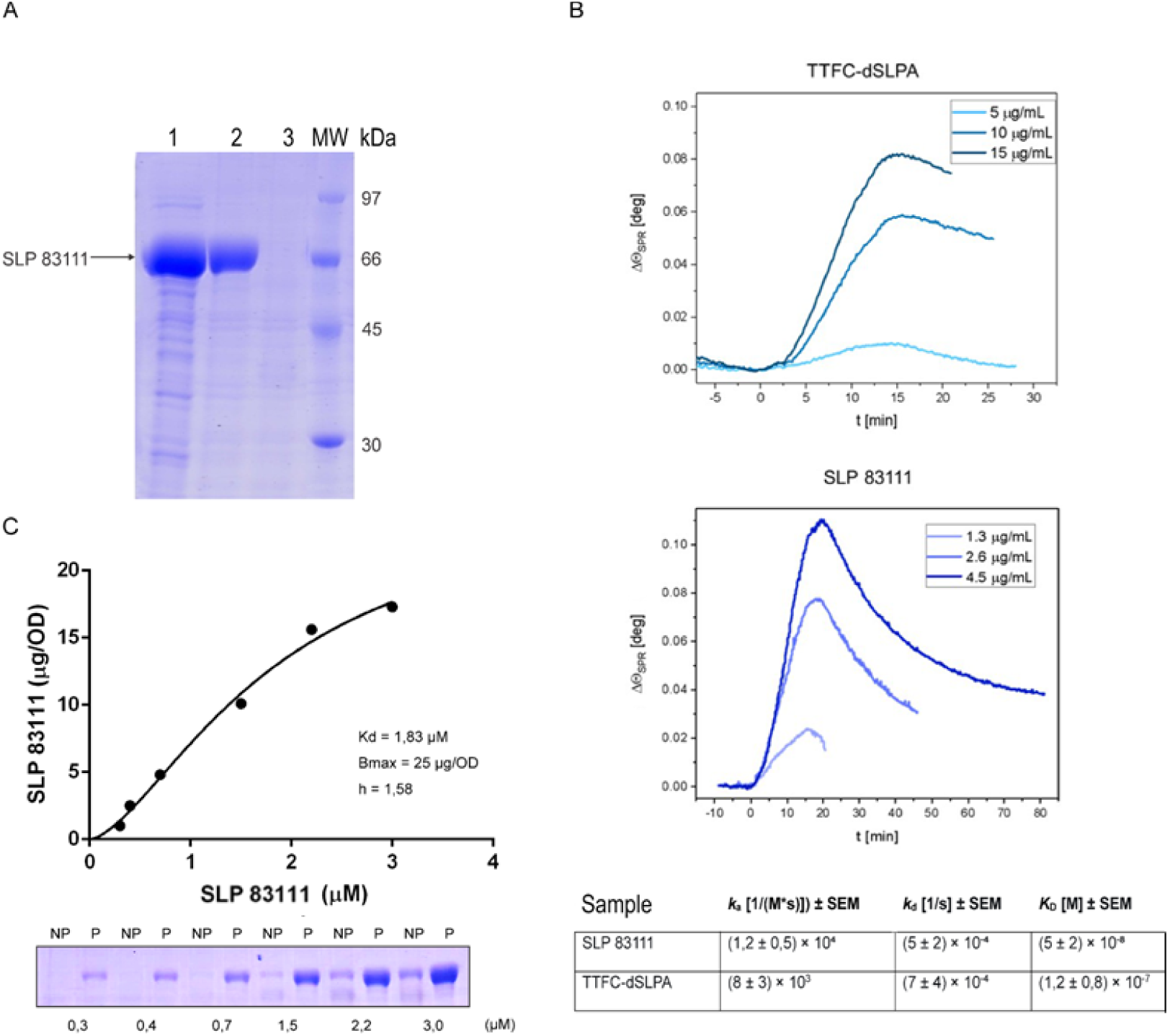
Production of native SLP 83111 and characterization of its interaction with the bacterial carrier. (A) SDS-PAGE analysis of SLP 83111 extraction from *L. kefiri* CIDCA 83111 by 5 M LiCl treatment. *L. kefiri* before extraction (lane 1), SLP 83111 extract (lane 2), *L. kefiri* after extraction (lane 3), molecular weight marker (lane 4). (B) MP-SPR analysis. Representative sensorgrams showing the change in the minimum SPR peak angular position (ΔΘ_SPR_) as a function of time during the interaction of each protein with the immobilized bacterial carrier. At t = 0 min the protein injection starts and at t = 10 min the PBS-T buffer flow is restored. Different analyte concentrations were measured under identical conditions, and k_a_, k_d_ and K_D_ were calculated from the average values obtained in independent fits (n = 3 independent experiments, four technical replicates each). Values are mean ± SEM. (C) Adsorption isotherm of SLP 83111 (upper panel) obtained using increasing protein concentrations (0.3-3 µM). The fraction bound to bacteria was quantified by densitometric analysis of SDS-PAGE (lower panel), protein bound to the carrier (P), unbound protein (NP). The dissociation constant K_d_ (1.83 µM) and the maximum binding capacity B_max_ (25 µg/OD of carrier) were determined by non-linear regression of the binding isotherm using GraphPad Prism 6.0.

The interaction of SLP 83111 and TTFC-dSLPA with the bacterial carrier was then characterized by two complementary approaches, MP-SPR to determine kinetic and affinity parameters and adsorption isotherms to assess the maximum binding capacity. Sensorgrams obtained by MP-SPR showed well-defined association and dissociation phases for both proteins, confirming their specific interaction with the immobilized carrier surface (Figure 1B). Kinetic analysis using a one-to-one interaction model yielded an association rate constant (k ), a dissociation rate constant (k_d_) and an equilibrium dissociation constant (K_D_) for each protein. SLP 83111 displayed high affinity for the bacterial surface, with a K_D_ in the 10^-8^ M range, indicating a stable and specific interaction. TTFC-dSLPA showed a slightly lower affinity (K_D_ = 1.2 x 10^-7^ M), consistent with previous observations for the interaction of TTFC-dSLPA with inactivated *B. subtilis* [13].

The binding capacity of native SLP 83111 to the bacterial carrier was further assessed by an adsorption isotherm. Keeping the amount of carrier constant and varying the concentration of SLP 83111, a maximum binding capacity (B_max_) of 25 µg/OD and an apparent dissociation constant (K_d_) of 1.83 µM were determined (Figure 1C). The apparent K_d_ obtained from the adsorption assay was approximately two orders of magnitude higher than the K_D_ determined by MP-SPR. This discrepancy is likely explained by the different nature of the two measurements: while MP-SPR monitors the interaction of purified protein with the immobilized bacterial surface in real time, the adsorption isotherm quantifies the amount of protein retained by whole bacteria after washing, where multivalent interactions, steric accessibility and surface heterogeneity may influence the apparent affinity. Despite this difference, both approaches indicate a strong and specific interaction that supports the stable immobilization of SLP 83111 on the carrier surface. Overall, these results confirm that both SLP 83111 and TTFC-dSLPA bind with high affinity to the bacterial carrier, a property that is relevant for the design of CBVs formulations co-displaying the antigen and the immunomodulator.

### 3.2 Co-incorporation of TTFC-dSLPA and SLP 83111 into CBVs

Once the interaction of both proteins with the carrier was characterized, the coating conditions were optimized to obtain CBVs formulations simultaneously displaying TTFC-dSLPA and SLP 83111. Two coating strategies were evaluated: simultaneous incubation of both proteins in the same mixture, and sequential coating, first with TTFC-dSLPA and then with SLP 83111. Sequential coating allowed a better co-incorporation of both proteins on the carrier (data not shown). Because SLP 83111 tended to displace TTFC-dSLPA when high concentrations were used, the amount of SLP 83111 was adjusted to achieve the simultaneous presence of both proteins. The optimal conditions for sequential coating were 1.8 µM TTFC-dSLPA followed by 1.4 µM SLP 83111.

The co-incorporation of TTFC-dSLPA and SLP 83111 on the bacterial carrier was confirmed by SDS-PAGE. As shown in Figure 2A, bands corresponding to TTFC-dSLPA (∼72 kDa) and SLP 83111 (∼71 kDa) were detected in the sequentially coated formulations. The presence of both bands in the bound fraction confirmed the co-immobilization of the two proteins on the carrier surface. To confirm the identity of the proteins in the different vaccine formulations, Western blot analysis was performed using specific polyclonal antisera against TTFC and SLP 83111. Detection with anti-SLP 83111 serum revealed a band of ∼71 kDa (native SLP 83111) in the CBVs-TTFC-SLP formulation (Figure 2B). Detection with anti-TTFC serum revealed a band of ∼72 kDa (TTFC-dSLPA) in the same formulation (Figure 2C). The absence of cross-reactivity between the two antisera confirmed the specificity of the detection. Together, these results validate the technical feasibility of co-formulating TTFC-dSLPA with SLP 83111 on the bacterial carrier and establish the sequential coating conditions that allow the simultaneous and stable presentation of both proteins.

**Figure 2.**
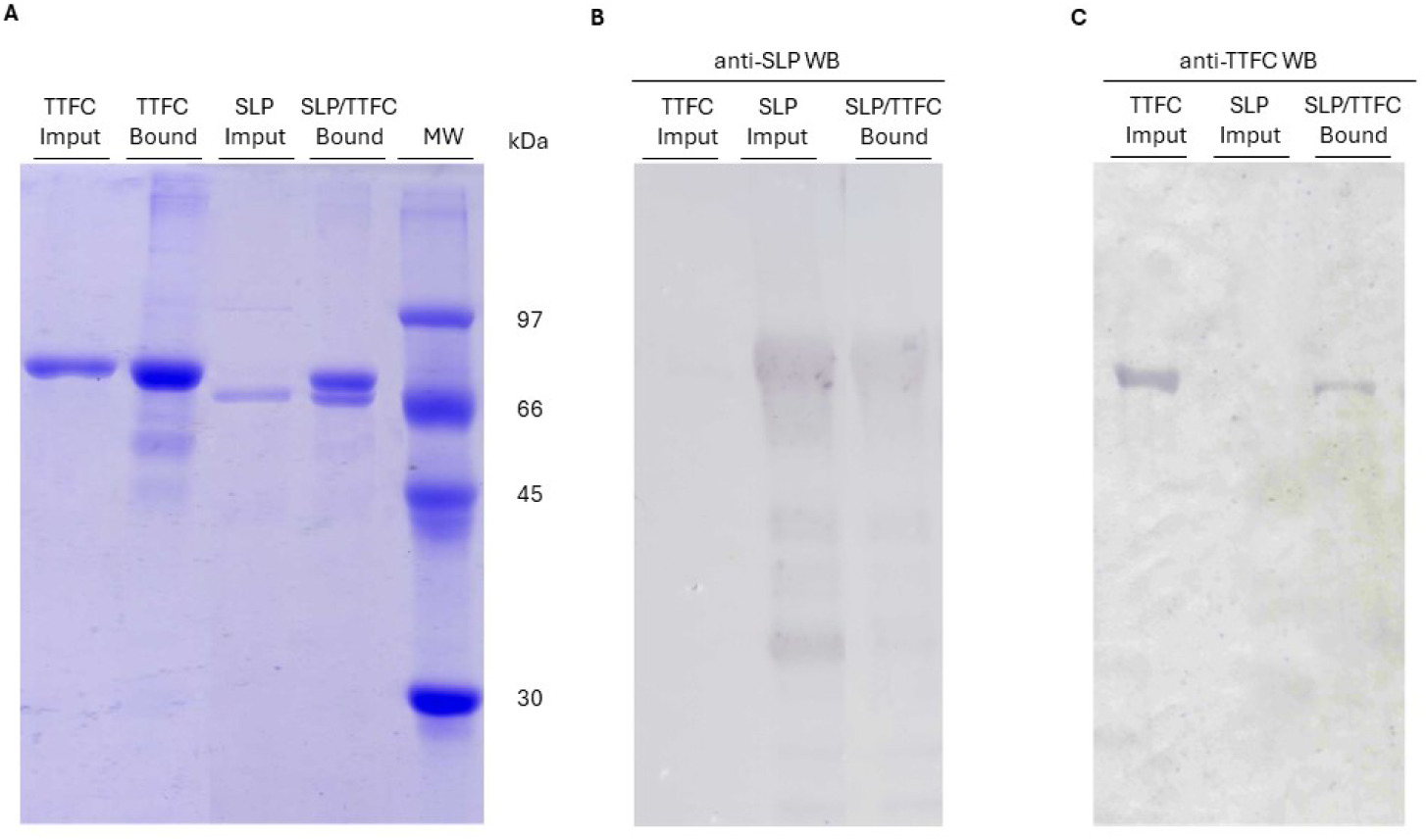
Characterization of CBVs formulations co-incorporating TTFC-dSLPA and SLP 83111. (A) SDS-PAGE analysis of CBVs-TTFC and CBVs-TTFC-SLP obtained by sequential coating. Purified TTFC-dSLPA input (lane 1), TTFC-dSLPA bound to the carrier (lane 2), purified SLP 83111 input (lane 3), TTFC and SLP 83111 simultaneously bound to the carrier surface (lane 4), molecular weight marker (MW, Low Molecular Weight Calibration Kit, lane 5). (B) Western blot analysis of the formulations revealed with anti-SLP 83111 serum or (C) with anti-TTFC serum. Purified TTFC-dSLPA input (lane 1), purified SLP 83111 input (lane 2), TTFC and SLP 83111 simultaneously bound to the carrier surface (lane 3).

### 3.3 SLP 83111 modulates the humoral response induced by CBVs

To evaluate the effect of SLP 83111 incorporation on the anti-TTFC humoral response induced by the CBVs platform, an immunization trial was performed in BALB/c mice using the three experimental vaccines described in Section 2.9 and summarized in Table 1. Total anti-TTFC IgG levels were first determined at 35 days post-immunization. EC_50_ IgG titers against TTFC-dSLPA for each group are shown in Figure 3A. Animals immunized with CBVs-TTFC reached the highest IgG titers (log_10_ 3.9). Mice receiving CBVs-TTFC-SLP showed a titer of log_10_ 3.5. Although anti-TTFC IgG levels in the CBVs-TTFC-SLP group were lower than those in the CBVs-TTFC group, they were significantly higher than those of the control group immunized with soluble TTFC-dSLPA (log_10_ 2.7). Figure 3B shows anti-TTFC IgG levels for individual sera (1:1,000 dilution). The overall behavior of each group reflected that observed with pooled sera. These results confirm that presentation of TTFC on the CBVs platform, regardless of SLP 83111 incorporation, induces an antibody response of greater magnitude than that obtained with the soluble antigen. The presence of SLP 83111 in the CBVs formulation did not substantially affect the magnitude of the total IgG response, although a slight decrease in titers was observed when the protein was incorporated.

**Figure 3.**
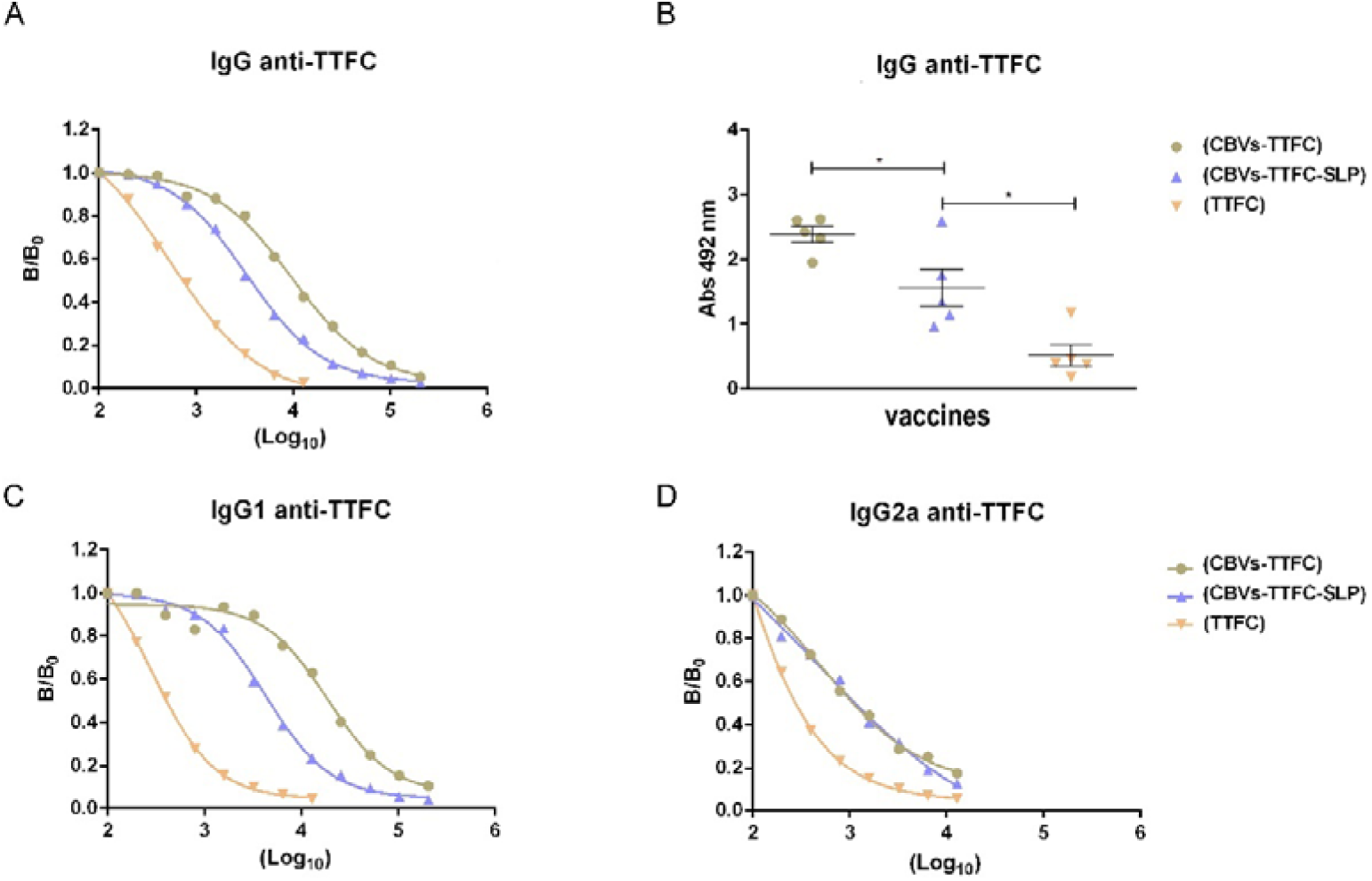
Humoral response induced by CBV formulations. (A) Total anti-TTFC IgG titers in serum. Serum samples were collected at 35 dpi and pooled. Anti-TTFC antibody titers were assessed by ELISA using serial dilutions, and EC_₅₀_ titers were calculated as the mid-log_₁₀_ of the reciprocal serum dilution. Titers for vaccines 1, 2 and 3 were 3.9, 3.5 and 2.7, respectively. (B) Total anti-TTFC IgG levels determined by ELISA at 35 dpi, expressed as absorbance values at 492 nm obtained from a 1:1,000 serum dilution. Data are mean ± SEM (n = 5/group). One-way ANOVA followed by Tukey’s multiple comparison test was used to analyze statistical significance between groups (*p < 0.05, **p < 0.01, ns, not significant). (C) Anti-TTFC IgG1 and (D) IgG2a antibody titers in serum. Serum samples were collected at 35 dpi and pooled. Titers were assessed by ELISA using serial dilutions and expressed as EC_₅₀_ values. Titers for vaccines 1, 2 and 3 were 4.2, 3.6 and 2.4 for IgG1, and 2.7, 2.9 and 2.4 for IgG2a, respectively. In A, C and D, pooled sera were used, and therefore no statistical analysis is shown.

To characterize the profile of the induced immune response, anti-TTFC IgG1 and IgG2a levels were determined in sera from immunized mice. EC_50_ IgG1 and IgG2a titers against TTFC-dSLPA for each group are shown in Figure 3C and 3D. Immunization with CBVs-TTFC induced a marked predominance of IgG1 over IgG2a (ratio 30:1) indicative of a Th2-polarized response, this result is consistent with previous observations for CBVs-TTFC and CBVs-GnRH [13, 14]. In contrast, incorporation of native SLP 83111 in the CBVs-TTFC-SLP formulation induced a more balanced response, with an IgG1/IgG2a ratio of 5:1. This shift was mainly due to a decrease in IgG1 titers (from log_10_ 4.2 to 3.6), whereas IgG2a titers remained essentially unchanged (from log_10_ 2.7 to 2.9). The control group immunized with soluble antigen showed very low levels of both isotypes, with an IgG1/IgG2a ratio of 1:1. Together, these results indicate that incorporation of native SLP 83111 into the CBVs-TTFC formulation modulates the humoral profile, reducing the Th2 bias and yielding a more balanced IgG1/IgG2a ratio.

### 3.4 Protection against lethal tetanus toxin challenge

To evaluate the protective capacity of the different formulations, immunized mice were challenged with 75 x LD_50_ of tetanus toxin (TeTx) on day 49 post-immunization. Survival and symptom onset were monitored for 10 days. As shown in Figure 4, mice immunized with CBVs-TTFC-SLP showed a delay in symptom onset and a higher survival rate than the other experimental groups. Although the difference between CBVs-TTFC and CBVs-TTFC-SLP was not statistically significant, a significant difference was observed between CBVs-TTFC-SLP and the soluble TTFC group, demonstrating the benefit of antigen presentation on the platform. Animals immunized with CBVs-TTFC showed intermediate survival, whereas the group immunized with soluble TTFC presented the lowest survival and the earliest onset of symptoms. These results indicate that incorporation of SLP 83111 into the CBV formulation not only modulates the humoral profile toward a more balanced IgG1/IgG2a ratio, but also tends to improve protection against tetanus toxin challenge, although the difference relative to CBVs-TTFC did not reach statistical significance under the conditions tested.

**Figure 4.**
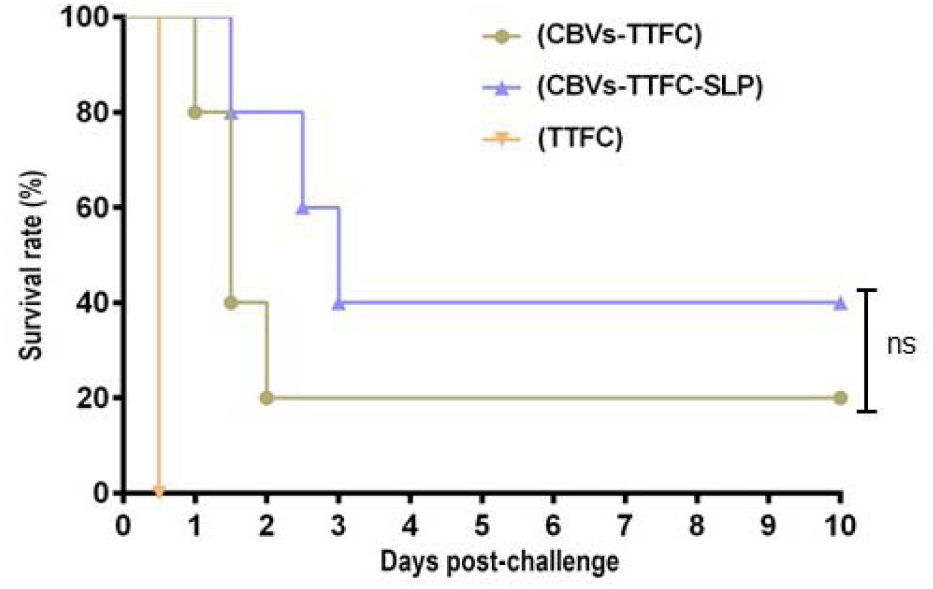
Survival curves of immunized mice after i.p. challenge with 75 × LD_₅₀_ of TeTx. Animals were monitored for 10 days post-challenge. Survival percentage is shown for each experimental group (n = 5/group). Statistical significance between survival curves was determined by the log-rank (Mantel–Cox) test. *p < 0.05 for CBVs-TTFC-SLP vs soluble TTFC; ns not significant for CBVs-TTFC vs CBVs-TTFC-SLP.

**Figure 5.**
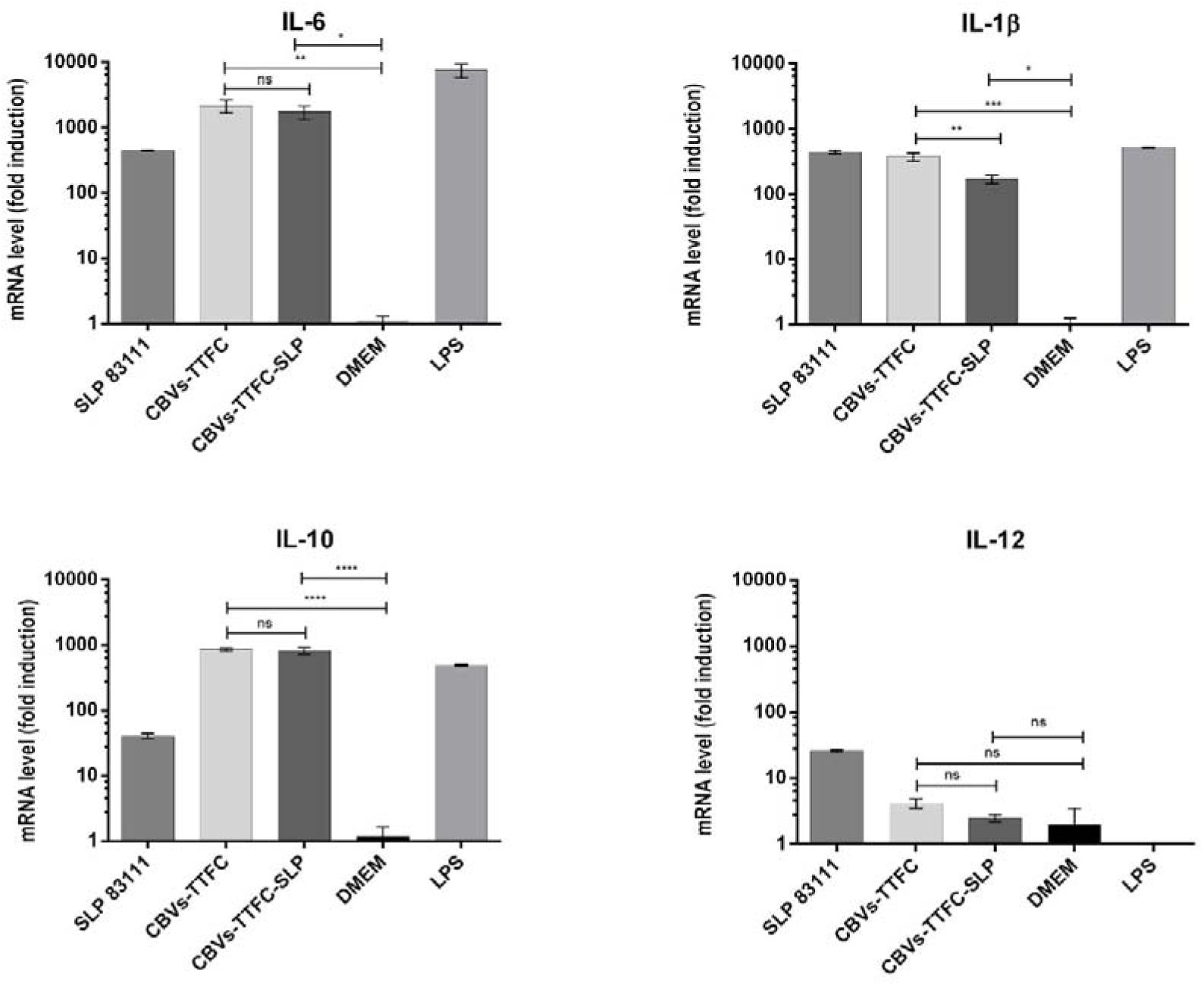
Cytokine expression in RAW 264.7 macrophages stimulated with CBV formulations. Quantification of IL-6, IL-1β, IL-10 and IL-12 mRNA by RT-qPCR after 6 h of stimulation with SLP 83111 (1 µg), CBVs-TTFC (2×10^7^ CBVs containing 2 µg of TTFC-dSLPA), CBVs-TTFC-SLP (2 ×10^7^ CBVs containing 2 µg of TTFC-dSLPA and 1 µg of SLP 83111), DMEM (negative control) or LPS (positive control). Data are expressed as mRNA induction (logarithmic scale) relative to the unstimulated control. Error bars indicate mean ± SEM of independent experiments (n = 3). Statistical significance was evaluated by one-way ANOVA followed by Tukey’s multiple comparison test (*p < 0.05, **p < 0.01, ***p < 0.001, ****p < 0.0001, ns; not significant).

### 3.5 SLP 83111 modulates cytokine expression in RAW 264.7 macrophages

To evaluate the effect of the formulations on macrophage polarization, mRNA levels of pro-inflammatory (M1) and regulatory (M2) cytokines were determined by RT-qPCR in RAW 264.7 macrophages after 6 h of stimulation with the different formulations. As shown in Figure 4, stimulation with CBVs-TTFC induced a significant increase in mRNA levels of the pro-inflammatory cytokines IL-6 and IL-1β compared with the unstimulated control (DMEM). When SLP 83111 was incorporated into the bacterial formulation (CBVs-TTFC-SLP), a differential modulation of the response was observed. IL-6 levels remained elevated and showed no significant differences between CBVs-TTFC and CBVs-TTFC-SLP. In contrast, IL-1β expression was significantly reduced in the CBVs-TTFC-SLP group compared with CBVs-TTFC.

IL-12 expression was low in all bacteria-stimulated groups, showing an almost complete suppression relative to the group treated with SLP 83111 alone. Notably, the positive control with LPS failed to induce IL-12 expression, consistent with the literature describing that the RAW 264.7 cell line has a deficient capacity to produce this cytokine. Regarding the regulatory cytokine IL-10, marked induction was observed both in the group stimulated with native SLP 83111 and in those treated with CBVs-TTFC and CBVs-TTFC-SLP. No significant differences were observed between the latter two groups, suggesting that CBVs themselves are a strong inducer of IL-10 and that addition of SLP 83111 does not interfere with this regulatory pathway. It should be noted that TNF-α determination, a classical M1 marker, is currently in progress and will be incorporated in future analyses. Overall, these preliminary results suggest that the CBV platform induces macrophage activation characterized by a mixed response, with strong production of IL-6 and IL-10, and that incorporation of native SLP 83111 specifically modulates the inflammatory response by attenuating IL-1β expression.

## 4. Discussion

The characterization of the interaction between SLP 83111 and the bacterial carrier was a prerequisite for the rational design of CBV formulations incorporating this immunomodulator. MP-SPR analysis showed that native SLP 83111 binds to inactivated *B. subtilis* with high affinity, with a K_D_ in the 10^-8^ M range. This value is lower-indicating stronger binding-than the K_D_ observed for TTFC-dSLPA (1.2 × 10^-7^ M) and considerably lower than the K_D_ reported for other S-layer-based anchoring systems, such as the SLAPTAG domain (4.7 x 10^-6^ M) [22]. The high affinity of SLP 83111 suggests that its interaction with cell wall components of *B. subtilis*, most likely teichoic and lipoteichoic acids [29], is stable under the experimental conditions tested. The adsorption isotherm additionally showed a maximum binding capacity of 25 µg/OD for SLP 83111. Although this value is lower than the 106 µg/OD determined for TTFC-dSLPA [13], it was sufficient for co-formulation with TTFC, allowing the simultaneous presentation of both proteins on the carrier.

The apparent K_d_ obtained from the adsorption isotherm (1.83 µM) was approximately two orders of magnitude higher than the K_D_ determined by MP-SPR. This difference is not unexpected and reflects the distinct nature of the two measurements. MP-SPR monitors the interaction of purified protein with an immobilized bacterial surface in real time and under flow, whereas the adsorption isotherm quantifies the amount of protein retained by whole bacteria after washing, a condition in which multivalent interactions, steric accessibility and surface heterogeneity can influence the apparent affinity. Similar discrepancies have been reported for other S-layer-cell wall interactions [30]. Importantly, both approaches indicate a strong and specific interaction, and together they support the conclusion that SLP 83111 can be stably displayed on the CBV surface alongside the antigen.

The most relevant finding of this work is the ability of native SLP 83111 to modulate the profile of the humoral response induced by the CBV platform. While CBVs-TTFC induced a marked predominance of IgG1 over IgG2a (ratio 30:1), characteristic of a Th2-polarized response, incorporation of SLP 83111 reduced this ratio to 5:1, indicating a more balanced profile. It is important to note that this shift was mainly driven by a decrease in IgG1 titers (from log₁₀ 4.2 to 3.6), whereas IgG2a titers remained essentially unchanged (from log₁₀ 2.7 to 2.9). Therefore, the effect of SLP 83111 should be interpreted as a reduction of the Th2 bias rather than as a switch toward a Th1 profile. This distinction is relevant because it implies that SLP 83111 does not simply redirect the response to the opposite pole, but rather moderates the strong Th2 polarization imposed by the intrinsic PAMPs of *B. subtilis*. Such a modulatory effect is consistent with the known ability of SLPs from *L. kefiri* to interact with C-type lectin receptors such as Mincle and SignR3, promoting dendritic cell maturation and cytokine secretion [16, 17], and with the observation that SLP 83111 can potentiate APC activation on its own [18].

The results of the tetanus toxin challenge showed that the CBVs-TTFC-SLP formulation tended to confer better protection than the other formulations, as evidenced by a delay in symptom onset and a numerically higher survival rate. However, the difference between CBVs-TTFC and CBVs-TTFC-SLP did not reach statistical significance under the conditions tested, and the only significant difference was observed between CBVs-TTFC-SLP and the soluble TTFC group. This result should be interpreted with caution, especially considering the limited sample size (n = 5 per group) and the inherent variability of survival experiments. Nevertheless, it is particularly interesting that the formulation with SLP 83111 showed slightly lower total anti-TTFC IgG titers than the formulation without SLP 83111 (log_10_ 3.5 vs 3.9), suggesting that the trend toward improved protection is not simply due to a higher magnitude of the antibody response but also to its quality. The more balanced IgG1/IgG2a profile could be associated with antibodies of higher neutralizing capacity or with a better activation of effector mechanisms dependent on antibody isotype, such as phagocytosis or antibody-dependent cellular cytotoxicity.

This observation is consistent with growing evidence indicating that cellular immunity should not be overlooked in protection against tetanus. Subcutaneous immunization with tetanus toxoid has been shown to induce a response with IgG1, IgG2a and IgG2b isotypes, indicative of a mixed Th1/Th2 response, and this response is associated with complete protection against a lethal challenge [31]. Similarly, immunization with recombinant *Lactobacillus plantarum* expressing TTFC induces IgG1 and IgG2b isotypes, and to a lesser extent IgG2a, also suggesting a mixed Th1/Th2 response [32]. Sublingual immunization with *B. subtilis* expressing TTFC has also been reported to promote a mixed Th1/Th2 response [33]. Together, these antecedents support the notion that a mixed Th1/Th2 response, rather than an exclusively Th2 polarization, is associated with more robust and optimal protection against tetanus. In the context of the CBV platform, the incorporation of native SLP 83111, by modulating the response toward a more balanced profile, could contribute to improving the functional quality of antibodies and, ultimately, the protection observed in this study.

In agreement with these in vivo findings, the in vitro stimulation of RAW 264.7 macrophages suggests that the CBV platform induces a mixed response at the cellular level. The high expression of IL-6 and IL-1β is consistent with pro-inflammatory (M1) activation, whereas the simultaneous production of IL-10 suggests the activation of regulatory pathways (M2a or M2c). The modulation observed with SLP 83111, evidenced by the reduction of IL-1β without altering IL-6 and IL-10 levels, suggests that this component is able to modulate the magnitude of the inflammatory response. Interestingly, IL-12 expression was low in all bacteria-stimulated groups, showing an almost complete suppression relative to the group treated with SLP 83111 alone. This observation is relevant because it indicates that the intrinsic PAMPs of *B. subtilis* dominate the cytokine profile when SLP 83111 is displayed on the bacterial surface. The mechanisms underlying this suppression remain to be elucidated, but several non-exclusive explanations can be proposed: (i) the density and spatial organization of *B. subtilis* PAMPs on the carrier surface may favor signaling pathways that suppress IL-12, such as those downstream of TLR2 [34]; (ii) the accessibility of SLP 83111 to its cognate lectin receptors may be reduced when the protein is anchored on the bacterial surface, limiting its ability to induce IL-12; and (iii) the internalization route of CBVs may differ from that of soluble SLP 83111, leading to different intracellular signaling outcomes. Further studies using receptor-blocking antibodies, confocal microscopy and cytokine profiling in dendritic cells will be required to address these hypotheses.

The capacity of SLP 83111 to modulate the Th1/Th2 balance of the response induced by CBVs is consistent with what has been reported for other S-layer proteins in different contexts. SlpA from *Lactobacillus helveticus* MIMLh5 induces the expression of TNF-α and cyclooxygenase 2 in human and murine macrophages, promoting a pro-inflammatory response in APCs [35]. The SLP from *Lactobacillus amylolyticus* NRIC 0558T induces IL-12 p40 secretion in the human monocytic cell line THP-1, suggesting a polarization toward Th1 [36]. Together with the results of the present work, these antecedents indicate that SLPs from different species and strains can exert diverse immunomodulatory effects depending on their structure and glycosylation. In the particular context of the CBVs platform, the incorporation of native SLP 83111 represents a novel strategy to modulate the immune response induced by the system. Unlike conventional adjuvants, which typically activate a limited signaling pathway and generate a predictable cytokine profile, SLP 83111 provides a diverse set of signals that could activate multiple PRRs simultaneously, generating a more complex immunological microenvironment and potentially more effective for the induction of robust and durable adaptive responses [6, 37].

Beyond the specific effect of SLP 83111, the most relevant finding of this work is that the CBVs platform itself can be rationally modulated through the incorporation of immunomodulatory molecules, extending its scope beyond antigen delivery. The platform was first shown to exert an adjuvant effect on a strongly immunogenic antigen (TTFC) [13] and was subsequently effective with a weakly immunogenic antigen (GnRH) [14]; the present study adds a third dimension by showing that the quality of the immune response can also be shaped. Together, these findings indicate that CBVs can be tailored not only to deliver antigens of diverse immunogenicity but also to direct the profile of the induced response, broadening the scope of the platform for the development of customized bacterial vaccines. This modularity is particularly attractive for veterinary applications, where the immune response requirements vary widely depending on the pathogen and the target species, and where the possibility of adjusting the formulation without modifying the antigen or the carrier represents a significant practical advantage.

This work has several limitations that should be acknowledged. First, the sample size in the challenge experiment (n = 5 per group) limits the statistical power to detect differences between CBVs-TTFC and CBVs-TTFC-SLP; larger studies will be needed to confirm whether the observed trend translates into a significant protection advantage. Second, the immunization experiment did not include control groups immunized with CBVs alone or with SLP 83111 alone, which would help to dissect the specific contribution of the immunomodulator versus the carrier. Third, the co-localization of TTFC-dSLPA and SLP 83111 was demonstrated at the level of the formulation (SDS-PAGE and Western blot), but not at the level of individual bacteria; immunofluorescence or flow cytometry with dual labeling would be required to formally demonstrate co-display on the same cell. Fourth, TNF-α determination, a classical M1 marker, was not available at the time of this study and will be incorporated in future analyses. Fifth, the doses of SLP 83111 used in the macrophage stimulation experiments (1 µg/well) were matched to those present in the CBVs-TTFC-SLP formulation, but the possibility that different doses might reveal additional effects cannot be excluded. Finally, the mechanism by which SLP 83111 modulates the response-particularly the suppression of IL-12 induction observed in stimulated macrophages remains to be fully elucidated.

## 5. Conclusion

The results of this work demonstrate that the CBVs platform can be rationally designed to modulate the immune response through the incorporation of the S-layer protein from *Lentilactobacillus kefiri* (SLP 83111) as an immunomodulator. The co-incorporation of native SLP 83111 with TTFC-dSLPA on the surface of inactivated *B. subtilis* yielded formulations in which both proteins were simultaneously displayed, and modified the profile of the humoral response, reducing the Th2 bias and shifting the IgG1/IgG2a ratio from 30:1 to 5:1. This modulation was accompanied by a trend toward improved protection against lethal tetanus toxin challenge, although the difference relative to the CBVs-TTFC formulation did not reach statistical significance; the significant advantage was observed relative to the soluble antigen, confirming the value of antigen presentation on the bacterial carrier.

From a platform-design perspective, the characterization of the interaction between SLP 83111 and the bacterial carrier provided the basis for co-formulation. The high affinity of SLP 83111 for the *B. subtilis surface* (K_D_ in the 10^-8^ M range), together with a maximum binding capacity of 25 µg/OD, ensures a stable and efficient presentation of the immunomodulator alongside the antigen. This property is essential for the development of modular vaccine platforms in which the co-presentation of antigens and immunomodulators can be controlled and adjusted according to need.

In conclusion, the results of this work, together with those obtained in our previous reports, position the CBVs platform as a versatile and robust tool for the development of next-generation vaccines. The ability to modulate the profile of the immune response through the incorporation of SLP 83111 opens new possibilities for the design of “tailored” vaccines adapted to the specific needs of each pathogen or application, and lays the foundation for future research aimed at exploring the application of the platform with other antigens of interest and other immunomodulatory molecules.

## Authors contribution statement

I.H. and M.H.A performed most of the experimental work, generated the figures, and significantly contributed to the writing and discussion of the article. M.S.H. conducted macrophage activation experiments. M.A.D.M. performed the multi-parametric surface plasmon resonance experiments. M.A.S participated in the writing. G.O.E., as the principal investigator and project director, contributed to the experimental design, interpretation of the results, and writing of the article.

## Ethics Statement

The experimental procedures of this study (protocol 010-42-24) were approved by the Institutional Animal Care and Use Committee of Exact Sciences Department at the National University of La Plata. Mice were immunized and challenged under the recommendations for animal experimentation (Helsinki Declaration and its amendments, and National Institutes of Health, United States: Guide for the Care and Use of Laboratory Animals).

